# Strategic stakeholders’ typology and mapping using stakeholder network analyses on integrated crops-livestock farming systems in West New Guinea

**DOI:** 10.1101/2020.07.06.189217

**Authors:** Deny Anjelus Iyai, Isti Widayati, Hendrik Fatem, Dwi Nurhayati, Maria Arim, Hanike Monim, Adolof Ronsumbre, Alnita Baaka, Lily Orisu, Desni T.R. Saragih, Yafet Syufi, Lambertus E. Nuhuyanan, Djonly Woran, Wolfram Mofu, Sangle Y. Randa, Lukas Y. Sonbait, Rizki Arizona, Michael Baransano, Daniel Seseray, Freddy Pattiselanno, Alexander Yaku, Johan Koibur

## Abstract

Stakeholders and its network play prominent roles in development particularly agriculture sector. The involvement of many stakeholders and other parties shaped how farms can sustain in terms of economic, social and environment indicators. Exploring the importance and roles of actors become strategic and vital to recognize. Study was done in Manokwari using focus group discussion towards twenty various represented individuals, groups and mass institutions. The queries discussed concerning background, resources delivery, interconnectivity amongst actors, intervention and innovation. The finding is that the stakeholders in mixed crop-livestock are dominated by individuals’ actors who privately manage the farms officially has laws. These actors are commonly act like stakeholders who are positively important ruled the farms. The threats are real and exist and should be lowering as much as possible to mitigate the turn-back effect. The top five shared resources are access, satisfaction, power, knowledge and time allocation. Those resources will stay longer to sustain strong needs of the farms. The relationship of actors is dominated by positive similarity and the ranges of correlation are varying in between negative, neutral to positive. This is due to actors reluctant to deliver the intervention and innovation. Actors with low interest and low power should then be promote to high interest and power by using aids, guidance and services from each actor in mixed crop-livestock farms business.

## INTRODUCTION

Agriculture development in particular crop-livestock sector is a mixed farming system that recognized and worked by many small-scale farmers in the world. The form of this farming is run by combining some commodities from crops and livestock. The trend of this system in the world is developed rapidly due to input efficiency, global climate changes and consumer concerns. These three reasons become the goal of sustainable development. In line with consumers concern, people now involve in determining products resulted from the farms. Development of this farming system is in fact done by involvement of many parties too.

Involvement of many parties such as individuals, groups and mass is to fulfill and satisfy people needs and consumers’ preferences. In Europe and other Western countries, crops and livestock products have been resulted from organic farms. Consumers and people now a days have been concerned about healthy food and food that produce without certain treatment. Caging animals in compartment are forbidden by animal welfare and right institution. Treat livestock with certain drugs and medicines are against the laws. The question raised now is what and who types of actors’ involvement, are they qualified and play vital rules in ensuring this policy of promoting animal right and welfare. Are these institutions already representing the consumers interest and answer the people and producers concerns.

Policies that ruled by the laws do not hamper once interest by legalizing other interest. This is done due to different perception how people see and perceive the objects. What constraint faced by mixed farming systems. Many publications of stakeholder and actor analyses discussed without seeing and analyzing the background and back-bound of the actors (Grimble & Wellard, 1997). Actors and stakeholders’ analyses commonly discussed qualitatively by drawing diagrams, pictures and connectivity lines. Whereas, many can be done by a bit more quantitatively compute the pattern and relationship of the network. Shapes of actors in line with individual, group and mass determine how actors have to be approached (Muniesa, 2015). Law status and types of organization become the criterion of legality in playing prominent roles (Hajjar *et al*., 2019). Legality will provide certainty and respect of involvement, beside trust worthy. Roles as stakeholder and shareholders will affect how contribution should be delivered in determining crop-livestock business beneficiary and production. Example is explained by Iyai *et al*. (2016) in Manokwari, West Papua-Indonesia.

Understanding the background and the back-bound of the actors are utmost important (Mayulu & Sutrisno, 2014). Best fitted and appropriate actors can play significant roles in promoting and sustaining cattle farming system particularly in Indonesia and specifically in West Papua. Iyai & Yaku (2015) identified several livestock farming systems in Manokwari, West Papua. Each livestock farming system established has certain relationship and typical involvement of various interest. Therefore, it is urgently needed to deeply digging up what characteristic of the institutions are, how it performs in real world livestock development. It is therefore needed to apply precise technical unit of analyses matched to predict the relationships of related and relevant stakeholders in benefiting economical- and social objectives of the crop-livestock farming systems. Characteristic of stakeholders or institutions can provide direction in executing implementing programs, aids, guidance and services in the near coming future.

One powerful social network analysis beside Gephi (Bastian *et al*., 2009), Netmap (Schiffer, 2007) and SmartPLS (Ringle *et al*., 2005), is Social Network Visualizer beside. The Social Network Analysis (SAN) is so far an adequate and appropriate software to compute network and relationship (Krupa *et al*., 2017). By mapping the stakeholders, institutions, which have no power and interest, would identify and in turn, will be easy to promote their roles comprehensively. This multi-sectors of agriculture development needs detail positioning of the roles and responsibilities from the involved actors. It is therefore, this study then aims to portrait typology of actors involved in old traditional livelihood of crop-livestock farming systems, i.e. mixed crop-livestock business based on West Papuan circumstances.

## METHODS

### Location and involved actors

Research was done in Manokwari, West Papua. We have chosen several organizations, groups and individuals who represented institutions, mass and households. We approached them using phone and invitation letter for collecting all relevant data and information concerning existing mixed crop-livestock farming business. Using focus group discussions and desk study from qualitative research (Moleong, 1991), relevant data collected consisted of information and data from research reports, policy documents, articles, daily newspapers and magazines. We considered doing this by the reasons that bunches of information and data written out and available even each was easy accessed.

We are concerned about the roles of stakeholders and shareholders in shaping and determining the development pattern of mixed crop-livestock business in West Papua, particularly in Manokwari. Manokwari was setup and developed as one of the central developments of mixed crop-livestock farms according to national plans of the Republic of Indonesia and by local livestock and veterinary provincial offices of West Papua province. All stakeholders grouped into local citizens, government, finance institutions (banks), markets, private and transportation.

### Data collection

During the research we collected information and data related to organizational function and characteristics of the mixed crop-livestock business-related stakeholders, i.e. shape of organization, status of low, types of organization, roles, effect and importance of organization. We also tried to collect data and information about threats and turn-back effect towards mixed crop-livestock farming development. In knowing the roles and presence of the stakeholders, we also recorded the sharing resources of organization, duration of period, continuity of the resources, power of resources and intervention done so far by organization.

### Method of analyses

In analyzing the power and flows of information amongst stakeholders, we used Social Network Visualizer (SocNetV). SocNetV (Kalamaras, 2019) is a cross-platform, light and free of charged social-stakeholder related software in network analyses and visualization. To visualize those graphs, we used PCC matrix, similarity matrix (SM), power centrality (PC), and Hierarchical clustering (HCA). The adjacency matrix of a social network (Supplement no. 1 & 2.) is a matrix where each element a(i,j) is equal to the weight of the arc from actor (node) i to actor j. If the actors are not connected, then a(i,j)=0. Computes the Cocitation matrix, C = A^T^ * A. C is an *n* x *n* symmetric matrix where each element (i,j) is the number of actors that have outbound ties/links to both actors i and j. The diagonal elements, C_ii_, of the Cocitation matrix are equal to the number of inbound edges of i (in Degree). A key notion in SNA is that of structural equivalence.

The idea is to map the relationships in a graph by creating classes or groups of actors who are equivalent in some sense. One way to do that, to identify groups of actors who are structurally equivalent, is to examine the relationships between them for similarity patterns. There are many methods to measure the similarity or dissimilarity of actors in a network. SocNetV supports the following methods: Similarity by measure and Pearson Correlation Coefficients. By applying one of these methods, SocNetV creates a pair-wise actor similarity/dissimilarity matrix. Computes a pair-wise actor similarity matrix, where each element (i,j) is the ratio of tie (or distance) matches of actors i and j to all other actors. In the case of Simple Matching, the similarity matrix depicts the ratios of exact matches of pairs of actors to all other actors. If the element (i,j) = 0.5, this means that actors i and j have the same ties present or absent to other actors 50% of the time. These measures of similarity are particularly useful when ties are binary (not valued). Computes a correlation matrix, where the elements are the Pearson correlation coefficients between pairs of actors in terms of their tie profiles or distances (in, out or both). The Pearson product-moment correlation coefficient (PPMCC or PCC or Pearson’s r) is a measure of the linear dependence/association between two variables X and Y. This correlation measure of similarity is particularly useful when ties are valued/weighted denoting strength, cost or probability. The Power Centrality (PC) is a generalized degree centrality measure suggested by Gil and Schmidt (1996a,b). For each node u, this index sums its degree (with weight 1), with the size of the 2nd-order neighborhood (with weight 2), and in general, with the size of the kth order neighborhood (with weight k). Thus, for each node u the most important other nodes are its immediate neighbors and then in decreasing importance the nodes of the 2nd-order neighborhood, 3rd-order neighborhood etc. For each node, the sum obtained is normalized by the total number of nodes in the same component minus 1. This index can be calculated in both graphs and digraphs but is usually best suited for undirected graphs. It can also be calculated in weighted graphs although the weight of each edge (u,v) in E is always considered to be 1 (therefore not considered). Hierarchical clustering (or hierarchical cluster analysis, HCA) is a method of cluster analysis which builds a hierarchy of clusters, based on their elements dissimilarity. In SNA context these clusters usually consist of network actors. This method takes the social network distance matrix as input and uses the Agglomerative “bottom up” approach where each actor starts in its own cluster (Level 0). In each subsequent Level, as we move up the clustering hierarchy, a pair of clusters are merged into a larger cluster, until all actors end up in the same cluster. To decide which clusters should be combined at each level, a measure of dissimilarity between sets of observations is required. This measure consists of a metric for the distance between actors i.e. Manhattan distance) and a linkage criterion (i.e. single-linkage clustering). This linkage criterion (essentially a definition of distance between clusters), differentiates between the different HCA methods. The result of Hierarchical Cluster Analysis is the clusters per level and a dendrogram. The concept of a clique in every life is pretty simple: a clique is a group of people who interact with each other much more regularly and intensely than with other people not belonging in the clique. That is, a group of people form a clique if they are all connected to each other. A clique is the largest subgroup of actors in the social network who are all directly connected to each other. In terms of graph theory, this notion is the same as a maximal complete subgraph of the equivalent graph of the social network. The word maximal means that for each clique the group of its members is expanded to include as many actors as possible; no other actors can be added to the clique. Essentially, a clique in Social Network Analysis consists of several overlapping closed triads.

SocNetV applies the Bron–Kerbosch algorithm to find all maximal cliques in an undirected or directed graph. It produces a census of all MAXIMAL cliques in the network and reports some useful statistics about these. The clique census report includes disaggregation by vertex and co-membership information. The Information Centrality (IC) is an index suggested by (Stephenson and Zalen, 1989) which focuses on how information might flow through many different paths. Unlike SC and BC, the IC metric uses all paths between actors weighted by strength of tie and distance.

The IC’ score is the standardized IC (IC divided by the sumIC) and can be seen as the proportion of total information flow that is controlled by each actor. Note that standard IC’ values sum to unity, unlike most other centrality measures. Since there is no known generalization of Stephenson & Zelen’s theory for information centrality to directional relations, the index should be calculated only for undirected graphs and is more meaningful in weighted graphs/networks. Note: to compute this index, SocNetV drops all isolated nodes and symmetrizes (if needed) the adjacency matrix even when the graph is directed Algorithm (Wasserman & Khaterine, 1994). In order to calculate the IC index of each actor, we create a N x N matrix A from the (symmetrized) sociomatrix with: Aii=1+di, Aij=1 if (i,j)=0, and Aij=1−wij if (i,j)=wij. Next, we compute the inverse matrix of A, for instance C, using the LU decomposition. Note that we can always compute C since the matrix A is always a diagonally strong matrix, hence it is always invertible. Finally, IC is computed by the formula: ICi−1Cii+T−2⋅RN, where: T is the trace of matrix C (the sum of diagonal elements) and R is the sum of the elements of any row (since all rows of C have the same sum). IC has a minimum value but not a maximum.

The steps in running this SocNetV version 2.5 presented in Figure 1. To catch the intervention shared by organization, we also look up into details what intervention done and shapes of innovation done by stakeholders. All data collectively typed into a Microsoft Excel worksheet and tabled into manuscript.

**Figure 1.**
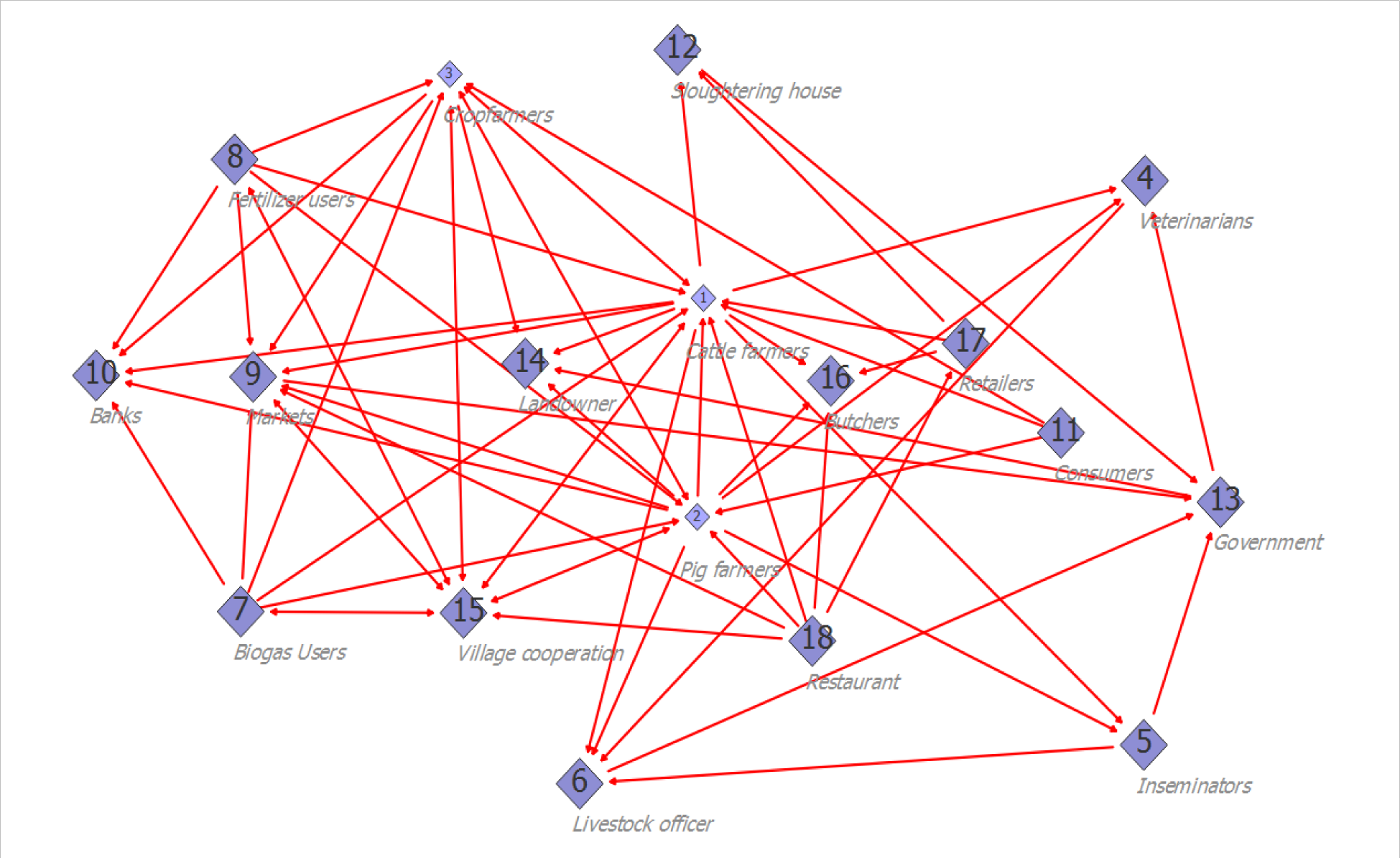
Mapping the involvement of actors amongst crop-livestock production systems.

## RESULTS

### Typology characteristic of organization

The recognized institutions or individuals who have been involved in determining and shaping mixed crop-livestock farming system and its business beneficiary are utmost important. Knowing the shapes of the organization status by law, types, roles, effect, importance, threats, and turn-back effect are seldom discussed by many authors. Shapes of organization as actors in leading crop-livestock farming systems grouped into three types, i.e. individuals (55.56%), group (38.89%) and mass (11.11%). We identified that the actors of mixed crop-livestock development ruled by law (50%) and the rest had no ruled by law. Types of organization established in mixed crop-livestock business sector were grouped in private and state institutions, subsequently 66.67% and 33.33%. The roles of organizations played by actors in crop-livestock farming systems were stakeholders (72.22%) and shareholders (27.78%).

Effects felt by goat business cycles on involved stakeholders were stated 12 actors had positive effect (66.67%) and only 8 actors in between had negative effect (44.47%). We interested in records the importance of the actors in ruled the crop-livestock business beneficiary. A number of 88.89% actors (16 organization) stated important and the rest had stated less important (22.11%). To assure the continuity of this business we measured the threat buried on business of cattle. We recorded 16 organizations had direct threat toward the development of crop-livestock production and the rest 2 actors had indirect effects. We finally eager to seek whether crop-livestock business beneficiary had turn-back effect amongst actors. The finding of this research reported no turn-back effect found inside 10 institutions (55.56%) and only 44.44% had turn-back effects. By knowing these fact characteristic of actors in reality, we concluded that cattle business beneficiary can sustain and has future development in West New Guinea.

### Available and status of resources

Shared resources inside crop-livestock business beneficiary cycles had some benefits, i.e. in the shapes of policy, finance, space, time, access, satisfaction, knowledge, skills, threat, power and feed materials. The finding and phenomenon faced by mixed crop-livestock farming systems was access (94.44%) and satisfaction in ranges of 83.33%. The shared resources can be offered in terms of power (66.67%), skills (61.11%), knowledge (55.56%), feed material (50%) and time (55.56%), space (44.44%), finance resources (44.44%) and lastly by policy (27.78%).

Duration of period in sharing resources organized by actors consisted of short term (22.22%) and long term periods (88.89%). Of actor profile, we found continuity of resources, i.e. sustain (50%) and unsustain (50%). Power of resources found was dominantly by strong power actors (50%), followed by weak power (27.78%) and neutral actors (22.22%). Weak power need further intervention and innovation in terms of resources’ needs. The need of Intervention was found in 9 actors (50%) and the rest were no need to intervene (50%). Delivery intervention can be made with related to policy, finance, knowledge, skills and relevant needs (Ventura *et al*., 2016). These types of intervention will further explain in the subsequent discussions.

To provide highlight of the position and how strength the relationship, we computed an analysis of stakeholder network analysis (SNA). The graph of Figure 2 highlighted the mental model of this relationship. The SNA output (Figure 3) depicted the picture of SNA based on Power centrality.

**Figure 2.**
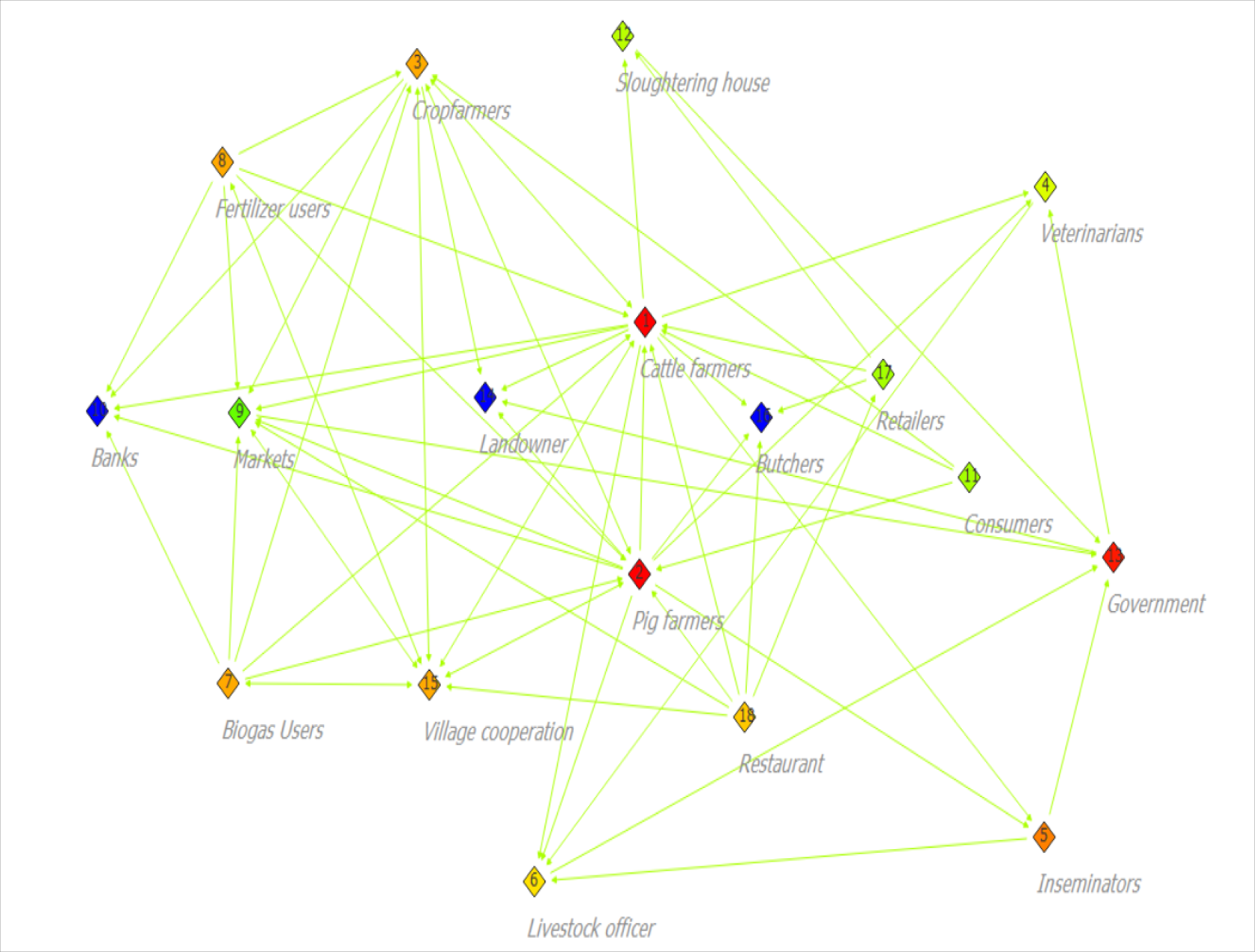
Stakeholder Network Analyses (SNA) of Cattle actors’ relationship based on Power centrality index and Kamada-Kawai (Force-directed model). Small and big size cubes indicated power relationship. Changed red to greed and blue colors indicating importance and strategic actors’ involvement from high to low power.

**Figure 3.**
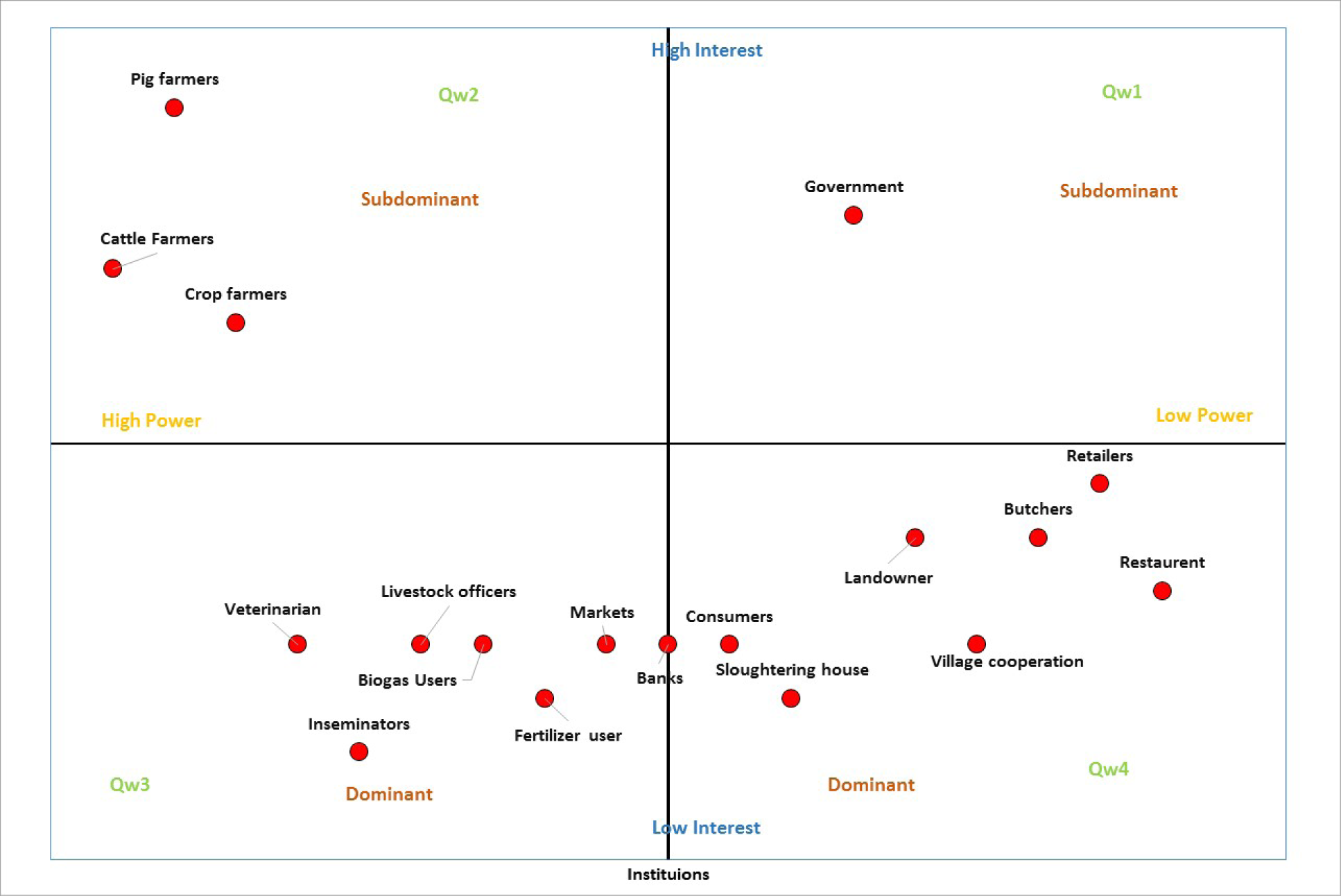
Stakeholder mapping on power and interest relationships under cattle farming systems

Of Figure 2 and Table 4., we succeeded in mapping interlinked relationship of actors’ network amongst crop-livestock farming in production systems. The output of SNA tell us the actors did not connect and the actors should have connection. Actors should connect are biogas users and fertilizer users, landowners and livestock officers, restaurant with government, village cooperation and government, retailers and village cooperation. Therefore, the responsibilities must be met by strategic and owner of policy makers, in this case government (central and provincial). In Central Java, constraints faced by mixed crop-livestock farmers made in causal loop diagram by Setianto *et al*. (2014).

**Table 1.** Stakeholders and roles and their responsibility under mixed crop-livestock development.

| No. | Institution | Role and Function |
| --- | --- | --- |
| 1. | Cattle farmers | Individuals and/or groups of farmers who are keeping cattle in their yards |
| 2. | Pig farmers | Individuals and/or groups of farmers who are keeping pigs in their yards |
| 3. | Crop farmers | Provide feed materials for men and animals |
| 4. | Veterinarian | Serving health of animals and farmers needs |
| 5. | Inseminators | Individuals who are serving the animal reproductions |
| 6. | Regency livestock offices | Ruled policy and regulation with related to cattle |
| 7. | Biogas users | Individuals who use gas as source of energy from livestock |
| 8. | Fertilizer user | Individuals who use fertilizer as source of organic soil materials from livestock |
| 9. | Market | Provide and distribute sale cuts |
| 10. | Banks | Providing saving account and loans |
| 11. | Consumer | Individuals who buy and consume the meat product |
| 12. | Slaughtering houses | Providing facilities for slaughtering livestock |
| 13. | Local government | Provide policy and regulations |
| 14. | Land owners | Providing width of areas for land use function |
| 15. | Village cooperation | Provide and distribute farmers need and production of farmers |
| 16. | Butchers | Individuals who do slaughter the meat of livestock |
| 17. | Retailers | Individuals and/or groups of community |
| 18. | Restaurants | Providing animal based product for consumers |

**Table 2.** Descriptive pattern of organization of actors in West New Guinea.

| No. | Typical institution | Sum | Proportion (%) |
| --- | --- | --- | --- |
| 1 | Shape of organization |  |  |
|  | Individual | 10 | 55.56 |
|  | Group | 7 | 38.89 |
|  | Mass | 2 | 11.11 |
| 2 | Law |  |  |
|  | Law | 9 | 50.00 |
|  | No law | 9 | 50.00 |
| 3 | Types |  |  |
|  | Private | 12 | 66.67 |
|  | State | 6 | 33.33 |
| 4 | Roles |  |  |
|  | Stakeholder | 13 | 72.22 |
|  | Shareholder | 5 | 27.78 |
| 5 | Effect |  |  |
|  | Positive | 12 | 66.67 |
|  | Negative | 8 | 44.44 |
| 6 | Importance |  |  |
|  | Important | 16 | 88.89 |
|  | Unimportant | 4 | 22.11 |
| 7 | Threat |  |  |
|  | Direct | 16 | 89.89 |
|  | Indirect | 2 | 11.11 |
| 8 | Turn-back Effect |  |  |
|  | Feedback | 10 | 55.56 |
|  | No feed-back | 8 | 44.44 |

**Table 3.**
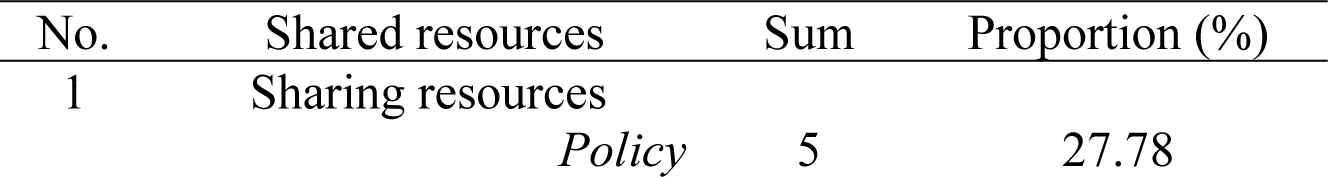
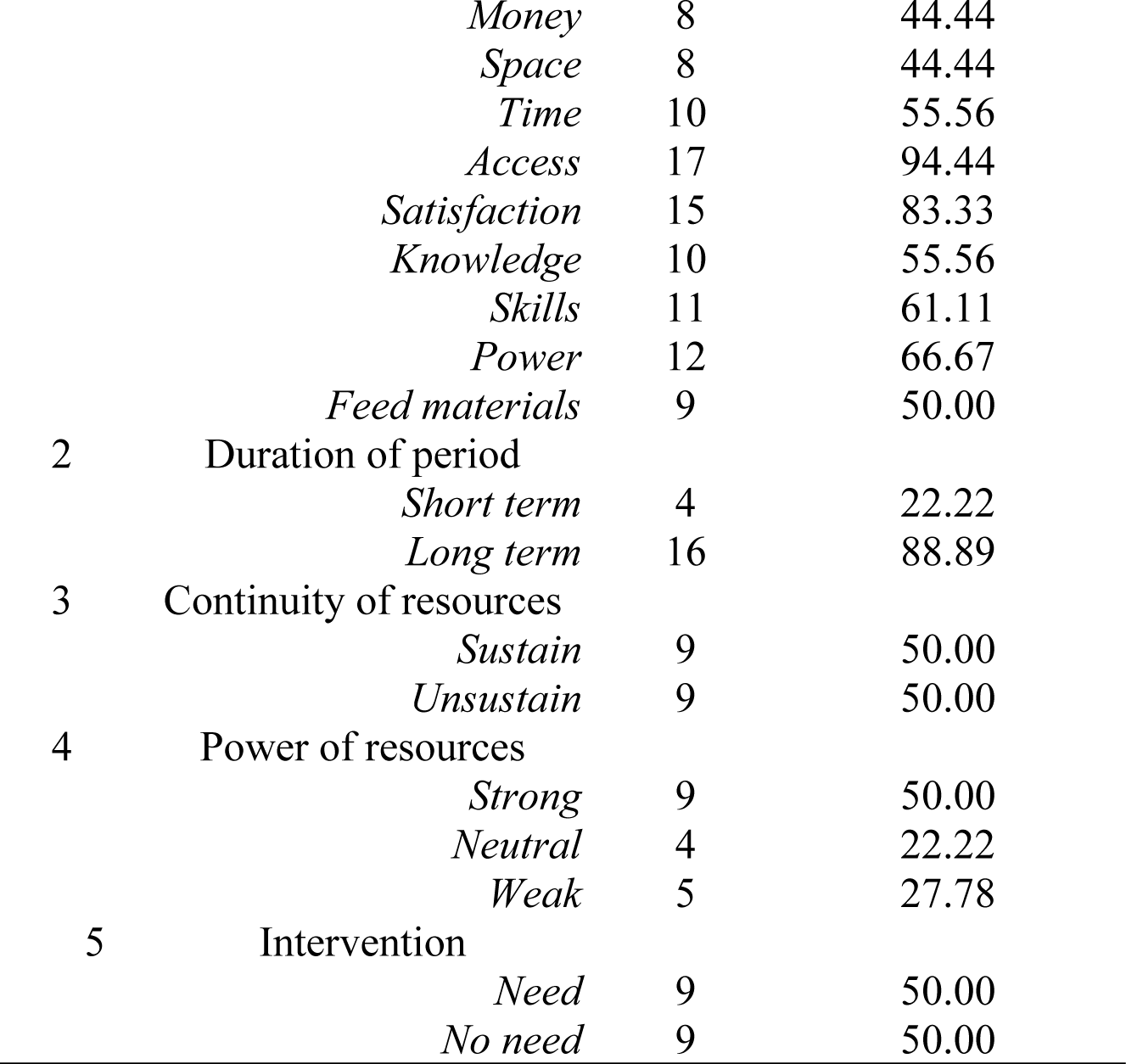
Identified shared resources of actors in West New Guinea

**Table 4.** Similarity Matrix; Matching coefficient (SMCC) of crop-livestock actors

| Actor\Actor | 1 | 2 | 3 | 4 | 5 | 6 | 7 | 8 | 9 | 10 | 11 | 12 | 13 | 14 | 15 | 16 | 17 | 18 |
| --- | --- | --- | --- | --- | --- | --- | --- | --- | --- | --- | --- | --- | --- | --- | --- | --- | --- | --- |
| 1 | 1.000 | 0.889 | 0.556 | 0.500 | 0.444 | 0.389 | 0.556 | 0.556 | 0.444 | 0.444 | 0.389 | 0.389 | 0.556 | 0.444 | 0.333 | 0.444 | 0.500 | 0.444 |
| 2 | 0.889 | 1.000 | 0.667 | 0.500 | 0.444 | 0.389 | 0.667 | 0.667 | 0.444 | 0.444 | 0.500 | 0.389 | 0.556 | 0.444 | 0.444 | 0.444 | 0.500 | 0.556 |
| 3 | 0.556 | 0.667 | 1.000 | 0.611 | 0.556 | 0.611 | 0.889 | 0.889 | 0.667 | 0.667 | 0.722 | 0.611 | 0.667 | 0.667 | 0.667 | 0.667 | 0.611 | 0.778 |
| 4 | 0.500 | 0.500 | 0.611 | 1.000 | 0.944 | 0.889 | 0.611 | 0.611 | 0.833 | 0.944 | 0.778 | 0.889 | 0.833 | 0.944 | 0.611 | 0.944 | 0.778 | 0.611 |
| 5 | 0.444 | 0.444 | 0.556 | 0.944 | 1.000 | 0.944 | 0.556 | 0.556 | 0.889 | 0.889 | 0.722 | 0.944 | 0.778 | 0.889 | 0.556 | 0.889 | 0.722 | 0.556 |
| 6 | 0.389 | 0.389 | 0.611 | 0.889 | 0.944 | 1.000 | 0.611 | 0.611 | 0.944 | 0.944 | 0.778 | 1.000 | 0.833 | 0.944 | 0.611 | 0.944 | 0.778 | 0.611 |
| 7 | 0.556 | 0.667 | 0.889 | 0.611 | 0.556 | 0.611 | 1.000 | 1.000 | 0.667 | 0.667 | 0.833 | 0.611 | 0.556 | 0.667 | 0.778 | 0.667 | 0.611 | 0.778 |
| 8 | 0.556 | 0.667 | 0.889 | 0.611 | 0.556 | 0.611 | 1.000 | 1.000 | 0.667 | 0.667 | 0.833 | 0.611 | 0.556 | 0.667 | 0.778 | 0.667 | 0.611 | 0.778 |
| 9 | 0.444 | 0.444 | 0.667 | 0.833 | 0.889 | 0.944 | 0.667 | 0.667 | 1.000 | 0.889 | 0.722 | 0.944 | 0.778 | 0.889 | 0.556 | 0.889 | 0.722 | 0.667 |
| 10 | 0.444 | 0.444 | 0.667 | 0.944 | 0.889 | 0.944 | 0.667 | 0.667 | 0.889 | 1.000 | 0.833 | 0.944 | 0.889 | 1.000 | 0.667 | 1.000 | 0.833 | 0.667 |
| 11 | 0.389 | 0.500 | 0.722 | 0.778 | 0.722 | 0.778 | 0.833 | 0.833 | 0.722 | 0.833 | 1.000 | 0.778 | 0.722 | 0.833 | 0.833 | 0.833 | 0.778 | 0.722 |
| 12 | 0.389 | 0.389 | 0.611 | 0.889 | 0.944 | 1.000 | 0.611 | 0.611 | 0.944 | 0.944 | 0.778 | 1.000 | 0.833 | 0.944 | 0.611 | 0.944 | 0.778 | 0.611 |
| 13 | 0.556 | 0.556 | 0.667 | 0.833 | 0.778 | 0.833 | 0.556 | 0.556 | 0.778 | 0.889 | 0.722 | 0.833 | 1.000 | 0.889 | 0.556 | 0.889 | 0.722 | 0.556 |
| 14 | 0.444 | 0.444 | 0.667 | 0.944 | 0.889 | 0.944 | 0.667 | 0.667 | 0.889 | 1.000 | 0.833 | 0.944 | 0.889 | 1.000 | 0.667 | 1.000 | 0.833 | 0.667 |
| 15 | 0.333 | 0.444 | 0.667 | 0.611 | 0.556 | 0.611 | 0.778 | 0.778 | 0.556 | 0.667 | 0.833 | 0.611 | 0.556 | 0.667 | 1.000 | 0.667 | 0.611 | 0.667 |
| 16 | 0.444 | 0.444 | 0.667 | 0.944 | 0.889 | 0.944 | 0.667 | 0.667 | 0.889 | 1.000 | 0.833 | 0.944 | 0.889 | 1.000 | 0.667 | 1.000 | 0.833 | 0.667 |
| 17 | 0.500 | 0.500 | 0.611 | 0.778 | 0.722 | 0.778 | 0.611 | 0.611 | 0.722 | 0.833 | 0.778 | 0.778 | 0.722 | 0.833 | 0.611 | 0.833 | 1.000 | 0.722 |
| 18 | 0.444 | 0.556 | 0.778 | 0.611 | 0.556 | 0.611 | 0.778 | 0.778 | 0.667 | 0.667 | 0.722 | 0.611 | 0.556 | 0.667 | 0.667 | 0.667 | 0.722 | 1.000 |

Down to Table 4., several actors 1^st^ to 18^th^ had positive clear similarity with SMCC= 0<C<1. Actors with SMCC=0 had no similarity at all. However, the value of SMCC>0, actors have same matches in their ties and/or distance. While SMCC=1 means the two actors have their ties to other actors exactly the same all the time. Actors in general had their SMCC>0 and found SMCC=1 for several relationships. Strong similarity seen in actors of regency livestock 6 vs slaughtering house 12 (C=1.000), followed by biogas users 7 vs fertilizer users 8; banks 10 vs land owners 14 and butchers 16. However, small SMCC also explained the strong and tied relationship. We encountered relationship of each actor to other actors and found dominancy of small SMCC. It tells that there is doubtful relationship amongst actors. The doubtful actors are cattle farmer1, pig farmers 2, biogas users 7, fertilizer users 8, village cooperation 15, and restaurant 18.

Down Table 5., several actors 1^st^ to 18^th^ had positive clear correlation with PCC=1.000 Actors with PCC=0.000 had no relationship at all. However, the rest had negative correlation (PCC<0.000). Actors had positive correlations were cattle farmers 1 vs pig farmers 2, crop farmers 3, veterinarian 4, biogas users 7, fertilizer users 8, local government 13, and retailers 17. Actors of pig farmers 2 had positive correlation with actor crop farmers 3, veterinarian 4, biogas users 7, fertilizer users 8, consumers 11, local government 13, village cooperation 15, retailers 17 and restaurants 18. Finally, actor 18 had positive correlation with actor pig farmers 2, crop farmers 3, biogas users 7, fertilizer users 8, market 9, consumers 11, village cooperation 15 and retailers 17.

**Table 5.** Matrix correlation coefficient of Pearson (PCC) of crop-livestock actors.

| Actor/Actur | 1 | 2 | 3 | 4 | 5 | 6 | 7 | 8 | 9 | 10 | 11 | 12 | 13 | 14 | 15 | 16 | 17 | 18 |
| --- | --- | --- | --- | --- | --- | --- | --- | --- | --- | --- | --- | --- | --- | --- | --- | --- | --- | --- |
| 1 | 1.000 | 0.878 | 0.323 | 0.228 | -0.048 | -0.293 | 0.244 | 0.244 | -0.048 | 0.000 | -0.098 | -0.293 | 0.293 | 0.000 | -0.221 | 0.000 | 0.293 | -0.035 |
| 2 | 0.878 | 1.000 | 0.595 | 0.228 | -0.048 | -0.293 | 0.522 | 0.522 | -0.048 | 0.000 | 0.293 | -0.333 | 0.293 | 0.000 | 0.051 | 0.000 | 0.041 | 0.244 |
| 3 | 0.323 | 0.595 | 1.000 | -0.200 | -0.293 | -0.200 | 0.870 | 0.870 | 0.153 | 0.000 | 0.488 | -0.200 | 0.098 | 0.000 | 0.418 | 0.000 | -0.041 | 0.467 |
| 4 | 0.228 | 0.228 | -0.200 | 1.000 | 0.683 | 0.000 | -0.200 | -0.200 | -0.098 | 0.000 | -0.124 | -0.067 | -0.067 | 0.000 | -0.200 | 0.000 | -0.124 | -0.200 |
| 5 | -0.048 | -0.048 | -0.293 | 0.683 | 1.000 | 1.000 | -0.293 | -0.293 | 0.429 | 0.000 | -0.182 | 0.683 | -0.098 | 0.000 | -0.293 | 0.000 | -0.182 | -0.293 |
| 6 | -0.293 | -0.293 | -0.200 | 0.000 | 1.000 | 1.000 | -0.200 | -0.200 | 0.683 | 0.000 | -0.124 | 1.000 | 0.000 | 0.000 | -0.200 | 0.000 | -0.124 | -0.200 |
| 7 | 0.244 | 0.522 | 0.870 | -0.200 | -0.293 | -0.200 | 1.000 | 1.000 | 0.153 | 0.000 | 0.620 | -0.200 | -0.293 | 0.000 | 0.709 | 0.000 | -0.041 | 0.467 |
| 8 | 0.244 | 0.522 | 0.870 | -0.200 | -0.293 | -0.200 | 1.000 | 1.000 | 0.153 | 0.000 | 0.620 | -0.200 | -0.293 | 0.000 | 0.709 | 0.000 | -0.041 | 0.467 |
| 9 | -0.048 | -0.048 | 0.153 | -0.098 | 0.429 | 0.683 | 0.153 | 0.153 | 1.000 | 0.000 | -0.182 | 0.683 | -0.098 | 0.000 | -0.174 | 0.000 | -0.182 | 0.153 |
| 10 | 0.000 | 0.000 | 0.000 | 0.000 | 0.000 | 0.000 | 0.000 | 0.000 | 0.000 | 0.000 | 0.000 | 0.000 | 0.000 | 0.000 | 0.000 | 0.000 | 0.000 | 0.000 |
| 11 | -0.098 | 0.293 | 0.488 | -0.124 | -0.182 | -0.124 | 0.620 | 0.620 | -0.182 | 0.000 | 1.000 | -0.124 | -0.182 | 0.000 | 0.620 | 0.000 | 0.179 | 0.289 |
| 12 | -0.293 | -0.333 | -0.200 | -0.067 | 0.683 | 1.000 | -0.200 | -0.200 | 0.683 | 0.000 | -0.124 | 1.000 | 0.000 | 0.000 | -0.200 | 0.000 | -0.098 | -0.200 |
| 13 | 0.293 | 0.293 | 0.098 | -0.067 | -0.098 | 0.000 | -0.293 | -0.293 | -0.098 | 0.000 | -0.182 | 0.000 | 1.000 | 0.000 | -0.293 | 0.000 | -0.182 | -0.293 |
| 14 | 0.000 | 0.000 | 0.000 | 0.000 | 0.000 | 0.000 | 0.000 | 0.000 | 0.000 | 0.000 | 0.000 | 0.000 | 0.000 | 0.000 | 0.000 | 0.000 | 0.000 | 0.000 |
| 15 | -0.221 | 0.051 | 0.418 | -0.200 | -0.293 | -0.200 | 0.709 | 0.709 | -0.174 | 0.000 | 0.620 | -0.200 | -0.293 | 0.000 | 1.000 | 0.000 | -0.041 | 0.313 |
| 16 | 0.000 | 0.000 | 0.000 | 0.000 | 0.000 | 0.000 | 0.000 | 0.000 | 0.000 | 0.000 | 0.000 | 0.000 | 0.000 | 0.000 | 0.000 | 0.000 | 0.000 | 0.000 |
| 17 | 0.293 | 0.041 | -0.041 | -0.124 | -0.182 | -0.124 | -0.041 | -0.041 | -0.182 | 0.000 | 0.179 | -0.098 | -0.182 | 0.000 | -0.041 | 0.000 | 1.000 | 0.367 |
| 18 | -0.035 | 0.244 | 0.467 | -0.200 | -0.293 | -0.200 | 0.467 | 0.467 | 0.153 | 0.000 | 0.289 | -0.200 | -0.293 | 0.000 | 0.313 | 0.000 | 0.367 | 1.000 |

Actor of cattle farmers had negative correlation with actor inseminators 5 (PCC=-0.048), regency livestock offices 6, market 9, consumers 11, slaughtering house 12, village cooperation 15, and restaurant 18 (PCC=-0.035). Actors had no correlation were cattle farmers 1 with banks 10, land owners 14 (PCC=0.000) and butchers 16 (PCC=0.000).

### Mapping interest and power

Down to Figure 3., it is interesting in mapping actors into other indicators of powers and interest. We considered this as importance due to organizational theoretical background (Grimble & Wellard, 1997). We grouped these two indicators into four quadrants (Qw1-Qw4). In the first quadrant (Qw1), we had government actor involved with low power and high interest. It is proven as well from Figure 2. that one of the red button is government showing the strategic and important actors, beside it has high interest. However, in the second quadrant (Qw2), we identified three actors, i.e. pig farmers, cattle farmers, and crop farmers, which had high power and high interest. Due to three actors, we consider as subdominant group.

Contrary with third quadrant (Qw3), seven actors were found and distributed in this quadrant. They apparently were actors with high power but had low interest as well. They were veterinarian, livestock officers, biogas users, inseminators, fertilizer users, market and banks. These actors dominantly distributed in this segment of relational roles and important players. The last segment is a fourth quadrant (Qw4) that was dominantly found filled by several actors. They were consumers, slaughtering houses, land owners, village cooperation, butchers, retailers, and restaurant.

Analyzing the places on quadrant by some actors, we suggest to promote several actors’ capacity building, roles and power. We aim to revitalize these organizations to have better roles and responsibility. Actors in the Qw1 (government) should move to the Qw2. Actors in the Qw3 (veterinarian, inseminators, livestock officers, biogas users, fertilizer users, markets and banks) should move as well in the Qw2. And finally, actors in Qw4 move to Qw2. This is done by reasons that actors will have better high interest and high power. Seeing this importation of actors’ network analyses (ANA), we pursued it by analyzing clustering using Hierarchical Cluster Analysis (HCA).

### Actors’ relationships

There were three leaves (Fig. 4.), i.e. simple (simplicifolius) consisted of actors 13 and 6, followed by double (bifolius) which consisted of actors 4 and 5, 9 and 15. And third one was triple (trifolius) which consisted of actor cattle farmers 1, pig farmers 2 and crop farmers 3; consumers 11, landowners 14, slaughtering houses 12; butchers 16, retailers 17 and restaurant 18; These had similarity in terms of roles and responsibility. The δ clade consisted of actor cattle farmers (1) and clade β which consisted of clades α (actors 2, 8, and 16) and actor 5.

**Figure 4.**
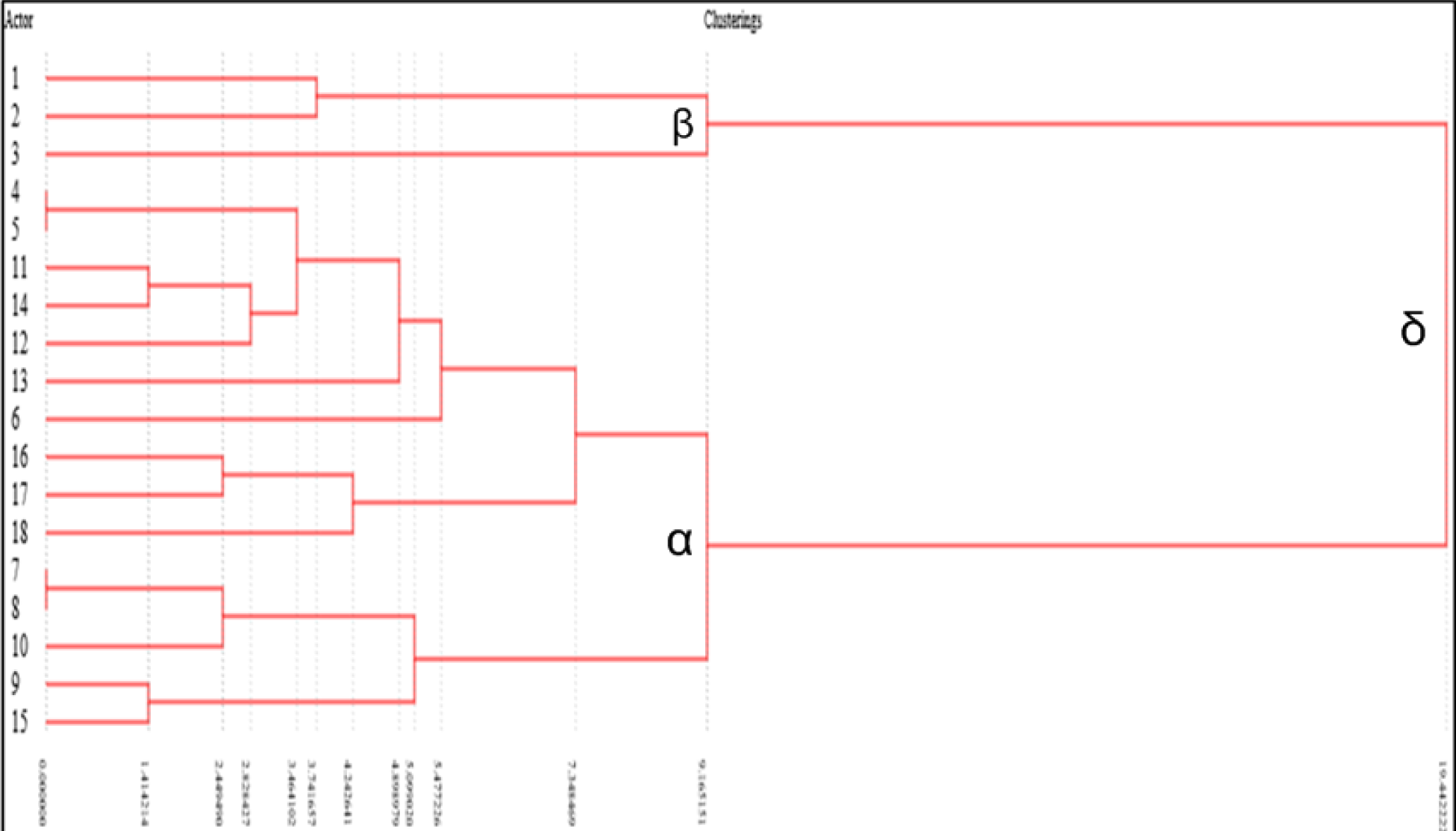
Hierarchical clustering analyses of crop-livestock actors’ relationship.

The δ clade consisted of actors β, i.e. cattle farmers 1, pig farmers 2 and cop farmers 3 and α which consisted of veterinarian 4, inseminators 5,…, village cooperation 15. Clades with similar height had similar to each other. Clades with dissimilar height had dissimilar relationship. Actors 4 and 5 along with actors 7 and 8 had a closed cluster relationship. The rests had far distance of cluster relationship.

### Intervention and Innovation

In assuring sustainability, intervention is utmost needs. We identified 11 actors needed policy intervention (61.11%). More than half 6 actors (33.33%) needed financial intervention. For instance, by improving grassland and/or pasture as reported by Oliveira *et al*., (2017). We found 4 else stakeholders which need spacing intervention (22.22%). Spacing intervention meant for infrastructure and wholesale cooperation, exampled in Thailand (Hasan *et al*. 2015). It seemed that no stakeholders needed intervention for time resource. In one hand more than 33.33% of actors (6) need access intervention. In few number of intervention of satisfaction was mentioned by an actor. Some actors (6) needed intervention of knowledge side (33.33%). Less than 38.89% (7 actors) needed intervention of skills. More than 33.33% of actors (6) needed intervention with related to threats they faced. Several actors (5) needed power intervention (27.78%), feed material (38.89%), and skills (38.89%), but some were requested for sustaining the cattle business beneficiary.

Differs from intervention, what innovations actually needed are questionable and shall be addressed to obtain clear concept and programs for improving crop-livestock business in West Papua. Innovation needs to assure the sustainability of crop-livestock farming systems. In policy sector, we found nine actors (50%) for performing policy innovation. Examples and experience reported by Gollnow & Lakes (2014). Specific innovation was regulation, law, standard operating procedures, research and development, monitoring and evaluation and taxation. Example explained by Hasan *et al*., (2015) in Makassar, Indonesia. In financial sector, four actors needed innovation of fund innovation followed by space for 10 actors (55.56%), six actors (33.33%) needed innovation for access. Satisfaction of actor services needed by 2 actors (11.11%), followed by innovation for knowledge needed by six actors (33.33%), skills (50%), power (5.56%) and feed materials (11.11%).

## DISCUSSIONS

Of Table 2, the typology of organization such as shapes, law status, types, roles, effect importance, even threat and turn-back effect will induce the rate and acceleration of each actor itself in establishing and delivering relationships and actions in mixed crop-livestock farming business. This portrait that mixed crop-livestock actors’ development in West New Guinea was on the stage of local and grass-root organization. National and International involved stakeholders are lagging behind for stimulating development. Experience so far shared generally by UNDP in almost region in West Papua and CIP-project in Wamena and Pegunungan Arfak. They have no bargaining position in determining the shapes and rate of crop-livestock development. The law of institutions determines the legality and power in sounding policy of development. Having access and trust for establishing cooperation and resources will induce acceleration development of mixed crop-livestock farming business. Distinguishing status of stakeholders and shareholders will enable easy-made and clear-contribution of delivering packages of the aids and services. Lowering negative effect in short run will enable actors to act with insurance. Direct threats are faced by many actors in crop-livestock farming system development. However, it then needs serious action in reducing direct impact. Sources of the threat are various, i.e. from animal health, wastes including livestock emission (Mariantonietta *et al*., 2017; Cardoso *et al*., 2016), forage management (Zanten *et al*., 2016) and price uncertainty (Asmarantaka *et al*., 2019). Internal and external warning should be addressed to avoid turn back effect.

Table 3. is inventorying possibilities of offered resources needed as inputs to stimulate development of crop-livestock farming system and enhancing farmer capacity including its actors. Eleven components of resources are found and therefore, it needs further policy and action to arrange it for establishing future and prospects of sustainable crop-livestock farming systems. Long term period shown how serious stakeholders in establishing livestock development. Even they can sustain and tend to have neutral and strong in pursuing targeted livestock development.

Table 4 grouped actors with similar typology and characteristic. These figures (2, 3 and 4) actually are drawing rich pictures and interpretation of actor network. We even have rich relationships and rich interlinked connectivity amongst actors. In Figure 2., various linking actors were created and these are phenomenal. It shows us the degree of mutual connectivity and as well as analyzing its prospect interlinked actors. Relationship between Table 2 and Table 3 along with Figure 2 and Figure 3 enable developing actors to be more precisely in delivering resources and capacities to share aids and guidance, added to this is service.

Table 5 explores the computed relational actors. It can be seen in Table 5 that, network and interlinked actors consist of positive, neutral and negative relationship. Meaning that negative network need adaptation and adjustment with local condition and targeted goals of crop-livestock development. Neutral relationship needs future intervention and innovation for driving its powers and interest in stimulating the tangible roles and future actions.

Table 6 investigated and recorded resources of further action can be done. Policy, skills and feed materials are the three top intervention that should deliver and needed by actors. However, according to Table 6 as well, policy, space and skill are the top three programs of innovation. Meaning that, actors shall bring and deliver intervention based on these priorities. In general, we convince the actors and/or donors and all et once convincing the receptors in promoting development of mixed crop-livestock farming business in West New Guinea, Indonesia.

**Table 6.**
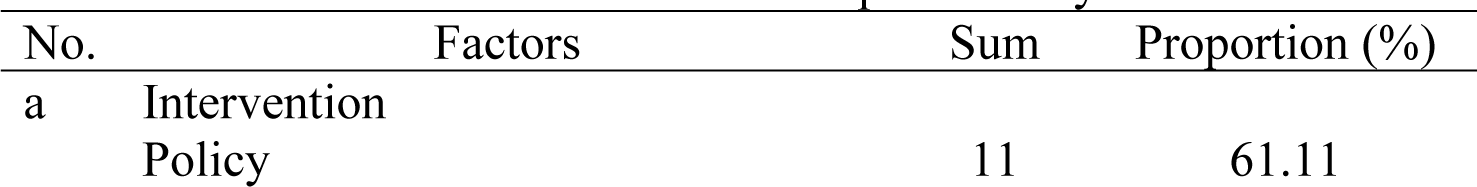
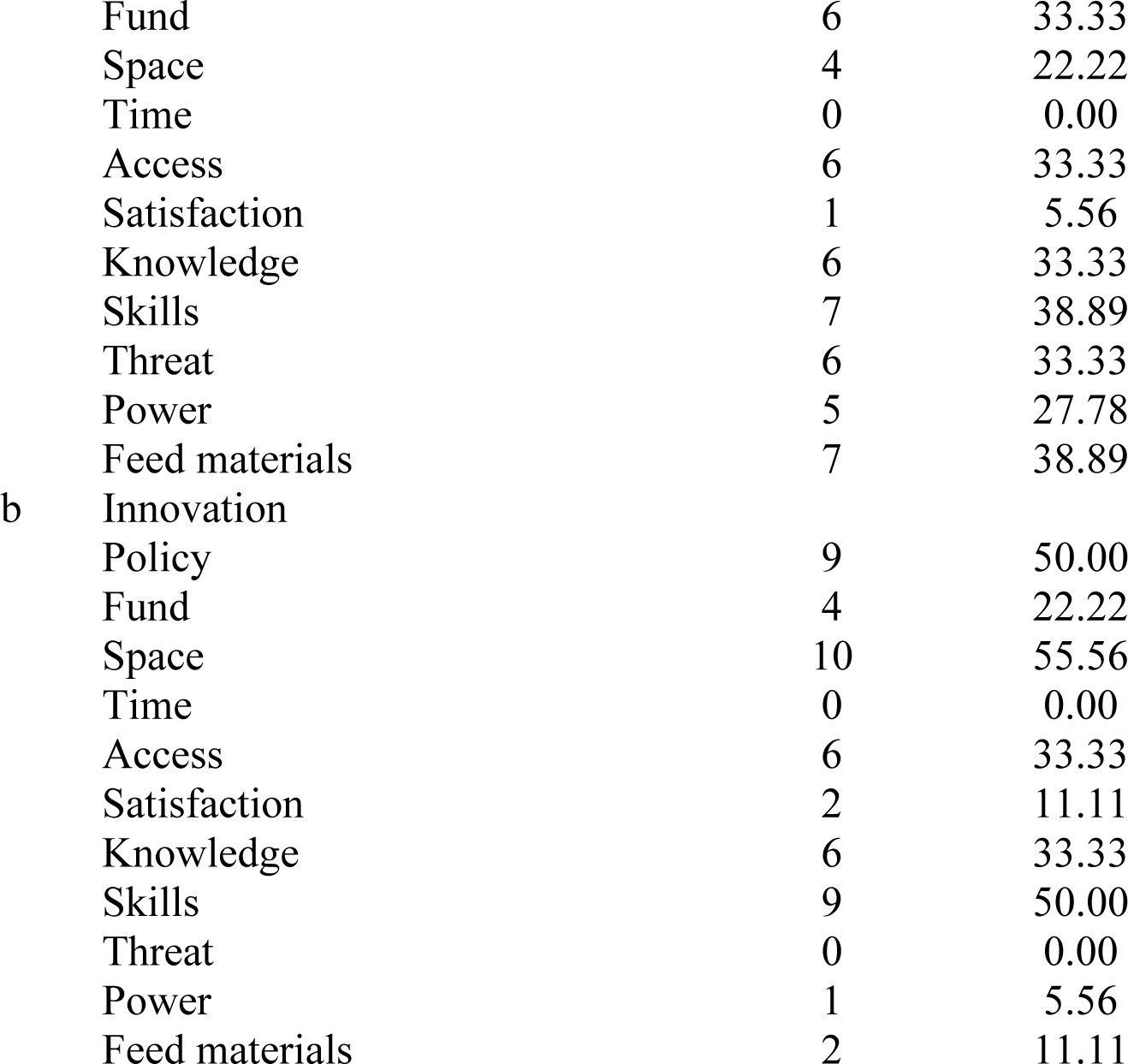
Intervention and innovation provided by cattle actors.

## CONCLUSIONS

We highlight the stakeholders in mixed crop-livestock are dominated by individuals’ actors who privately manage the farms officially has laws. These actors are commonly act like stakeholders who are positively important ruled the farms. The threats are real and exist and should be lowering as much as possible to mitigate the turn-back effect. The top five shared resources are access, satisfaction, power, knowledge and time allocation. Those resources will stay longer to sustain strong needs of the farms. The relationship of actors is dominated by positive similarity and the ranges of correlation are varying in between negative, neutral to positive. This is due to actors reluctant to deliver the intervention and innovation. Actors with low interest and low power should then be promoted to high interest and power by using aids, guidance and services from each actors in mixed crop-livestock farms business.

## ACKNOWLEDGEMENT

Authors wish to thank all represented institutions both states and/or private even individuals in sharing experience, information and data.

